# deepTFBS: Improving within- and cross-species prediction of transcription factor binding using deep multi-task and transfer learning

**DOI:** 10.1101/2025.03.19.644233

**Authors:** Jingjing Zhai, Yuzhou Zhang, Chujun Zhang, Xiaotong Yi, Minggui Song, Chenglong Tang, Pengjun Ding, Zenglin Li, Chuang Ma

**Author notes:** **Corresponding Author** Chuang Ma, State Key Laboratory of Crop Stress Resistance and High-Efficiency Production, Center of Bioinformatics, College of Life Sciences, Northwest A&F University, Shaanxi, Yangling 712100, China. These authors contribute equally to this work (Jingjing Zhai, Yuzhou Zhang).

## Abstract

The precise prediction of transcription factor binding sites (TFBSs) is crucial in understanding gene regulation. In this study, we present deepTFBS, a comprehensive deep learning (DL) framework that builds a robust DNA language model of TF binding grammar for accurately predicting TFBSs within and across plant species. Taking advantages of multi-task DL and transfer learning, deepTFBS is capable of leveraging the knowledge learned from large-scale TF binding profiles to enhance the prediction of TFBSs under small-sample training and cross-species prediction tasks. When tested using available information on 359 *Arabidopsis* TFs, deepTFBS outperformed previously described prediction strategies, including position weight matrix, deepSEA and DanQ, with a 244.49%, 49.15%, and 23.32% improvement of the area under the precision-recall curve (PRAUC), respectively. Further cross-species prediction of TFBS in wheat showed that deepTFBS yielded a significant PRAUC improvement of 30.6% over these three baseline models. deepTFBS can also utilize information from gene conservation and binding motifs, enabling efficient TFBS prediction in species where experimental data availability is limited. A case study, focusing on the *WUSCHEL* (*WUS*) transcription factor, illustrated the potential use of deepTFBS in cross-species applications, in our example between *Arabidopsis* and wheat. deepTFBS is publically available at https://github.com/cma2015/deepTFBS.

**One sentence summary:** The high-performing deep learning framework deepTFBS can flexibly perform within- and cross-species prediction of TF binding, transferring the transcription regulation from model to non-model species.

## INTRODUCTION

The complex regulation of gene expression, governed by *cis*-regulatory elements (CREs) and trans-acting factors in eukaryotes, enables cells to achieve precise responses during different developmental stages or under various environmental conditions. Transcription factors (TFs) are key trans-factors that regulate gene expression by recognizing specific DNA sequences, TF binding sites (TFBSs) [1, 2]. Various high-throughput experimental methods have been developed to determine the DNA sequence specificity of TFs. These include the chromatin immunoprecipitation (ChIP) assay with sequencing (ChIP-Seq) *in vivo* [3] and DNA affinity purification sequencing (DAP-Seq) *in vitro* [4, 5]. Utilizing these techniques, genome-wide TFBS profiling experiments have been conducted in multiple plant species, including *Arabidopsis thaliana* [4, 6], *Zea mays* [7, 8], *Triticum aestivum* [9] and *Triticum Urartu* [10]. These work generated valuable resources and provided considerable insight into the complex TF-target regulation involved in various biological processes in plants. However, such work is costly, labor-intensive, and time-consuming, limiting the number of species in which TFBSs can be experimentally mapped. Therefore, there is an urgent need for computational methods to identify the syntax of TF-DNA interactions based on available resources in order to apply these to other organisms, where the availability of experimental datasets is limited.

Numerous computational algorithms have been developed to identify the binding preferences of TFs within a single species. The most commonly used method is the position weight matrix (PWM), which uses independent positional statistics of four nucleotides (A, C, G and T) to predict TFBSs by scanning the DNA sequences of interest. However PWM ignores the weak signal and contextual information of TF binding patterns, often leading to false positive binding site predictions and adversely affecting downstream analyses [11]. In contrast, machine learning (ML) approaches, particularly recently developed deep learning (DL) algorithms have emerged as powerful alternatives. These advanced computational methods can effectively learn complex patterns, including long-distance nucleotide dependencies, by leveraging large-scale training data [12–14]. ML/DL algorithms have led to improvements in TFBS prediction accuracy using DNA sequences alone [15–23]. Beyond sequence data, these approaches have also successfully incorporated various DNA structural features, including DNA shape [22, 24, 25], and chemical and structural properties [26]. Among the DL-based methods, DeepBind [15] first utilized a convolutional neural network (CNN) to model DNA and RNA binding specificities, achieving superior prediction accuracy with both *in vitro* and *in vivo* data. Subsequently, more sophisticated DL models were developed for predicting DNA regulatory elements in human, such as DeepSEA [17], DanQ [18] and BPNet [23]. While DeepSEA utilizes a CNN architecture to learn the regulatory patterns from DNA sequences, DanQ implements a hybrid approach combining CNN with recurrent neural networks to enhance prediction capabilities. BPNet was specially desinged to predict base-resolution ChIP-nexus profiles and provide insights into TF binding patterns and their interactions. More recently, transformer-based models such as BERT-TFBS [21] have emerged, leveraging the power of attention mechanisms to capture complex dependencies in DNA sequences and improve TFBS prediction accuracy. Despite these advances in DL-based approaches, several key challenges remain to be addressed.

First, DL-based methods typically rely on substantial amounts of training data to achieve their accuracy. When these methods are applied in prediction tasks where experimentally reported binding data is limited, their prediction accuracy decreases markedly [25, 27]. In addition, the existing DL-based methods do not always perform well when used in cross-species TFBS prediction, even when the TFs are highly conserved or represent direct orthologs. This limitation is partly due to the high turnover of TF binding sites [28]. In human and mouse, domain-adaptive neural network [29] and adversarial training [30] have been explored to enhance the cross-species prediction of binding sites. However, few studies have explored effective ML/DL strategies for cross-species TFBS prediction in plants. Considering the scarcity of experimental TF binding data in many plant species, the development of algorithms suitable for the prediction of TFBS within limited datasets remains of great importance and achieving accuracy in cross-species use would be an extremely important goal.

This study introduces deepTFBS, a DL framework designed for the precise identification of TFBSs within and across species. By leveraging multi-task and transfer learning, deepTFBS effectively overcomes the limitations associated with small training sample sets in building predictive models. In addition, for cross-species predictions, we propose the integration of orthologous targets and PWM information to enhance accuracy. The utility of deepTFBS was validated using extensive publicly accessible TF binding profile data in *Arabidopsis* and wheat. Moreover, using the *WUS* TF as a test case, we experimentally confirmed several predicted binding events in wheat, using yeast one-hybrid assays, further demonstrating the accuracy of TFBS predictions provided by deepTFBS.

## RESULTS

### Designing the deepTFBS framework

deepTFBS is a DL framework specifically designed to accurately predict TFBSs, incorporating multi-task (deepTFBS-MT) and transfer learning (deepTFBS-TL) strategies. In training phase I, the multi-task DL model deepTFBS-MT is constructed by simultaneously learning multiple TF binding preferences (**Figure 1A**). deepTFBS-MT is pre-trained on binding data from 359 *Arabidopsis* TFs. It processes input sequences of 1,000 base pairs (bp), encoded using one-hot encoding (see METHODS). These encoded sequences are fed into the deepTFBS backbone, which constitutes the core network architecture of the framework (**Figure 1B**). The model outputs the binding probabilities for all 359 TFs via a fully connected layer comprising 359 neurons, each corresponding to a specific TF (**Figure 1A**). The pre-trained deepTFBS-MT offers two key advantages. Firstly, by fitting a single model with all available TF binding profiles, deepTFBS-MT leverages a large dataset to train an accurate, robust, and generalizable DL model. Second, it utilizes on the strengths of multi-task learning to deliver more precise and unbiased TFBS predictions.

**Figure 1.**
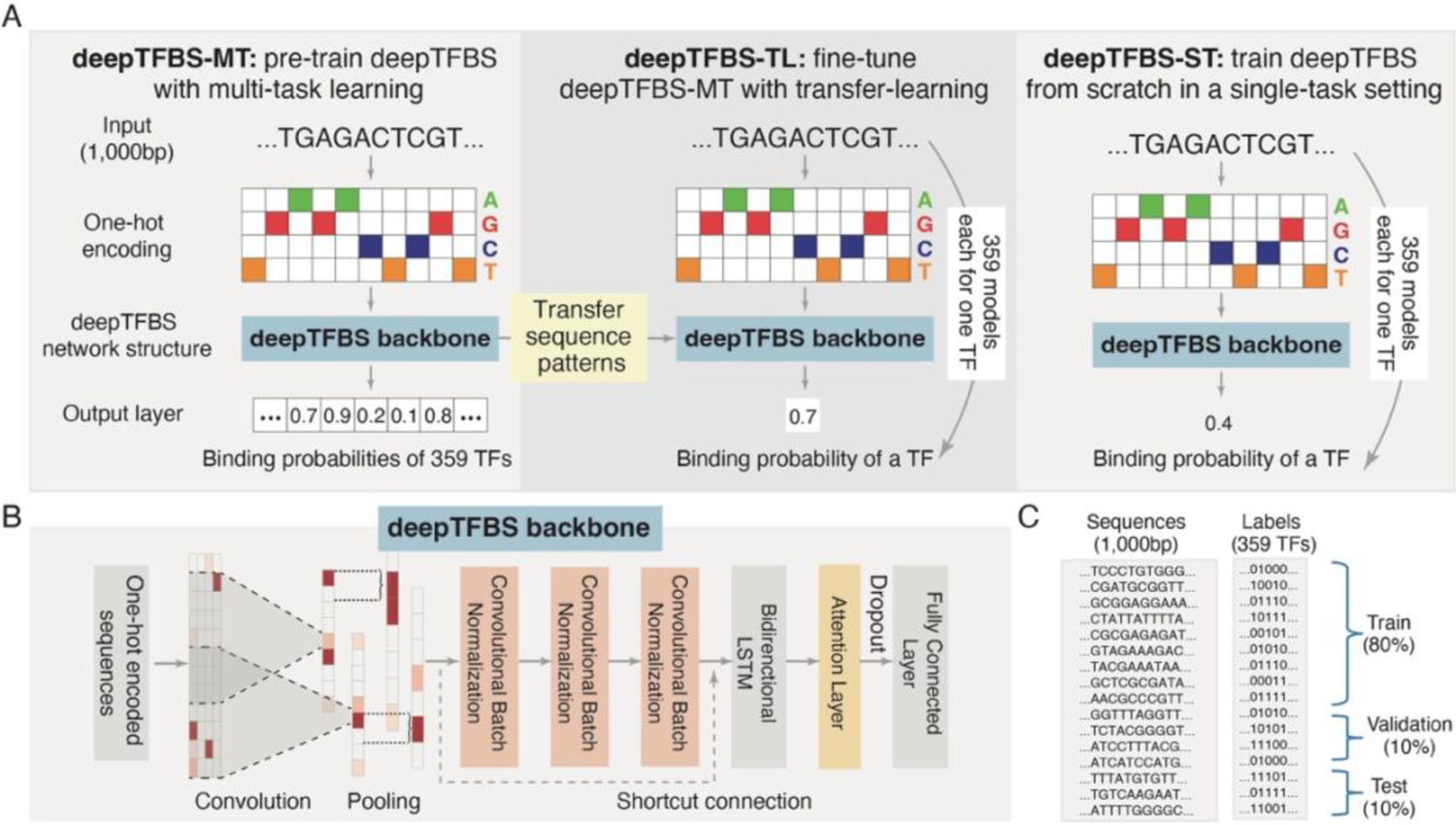
Illustration of deepTFBS framework. (A) The three variants of the deepTFBS framework. (B) The deepTFBS network structure backbone. (C) Train, validation and test data splits in deepTFBS-MT.

However, training deepTFBS-MT requires large-scale TF binding profiles, to address the challenge of limited TF binding profiles in certain species, we developed deepTFBS-TL, a transfer learning (TL)-based method that builds upon the foundation of deepTFBS-MT (**Figure 1A**). In this implementation, a binary classification model is trained for each TF, utilizing the sequence patterns learned by deepTFBS-MT. Specifically, deepTFBS-TL initializes the model with weights learned from deepTFBS-MT, effectively narrowing the parameter search space. Then, deepTFBS-TL uses a new fully connected layer to replace the output layer for binary classification. The transfer learning technology used is particularly advantageous for model training with small datasets. For comparison, we also implemented deepTFBS-ST, which trains a similar binary classification model but without transfer learning (starting from randomly initialized weights). This serves as a baseline to evaluate the benefits of transfer learning in deepTFBS-TL (**Figure 1A**).

Both deepTFBS-MT and deepTFBS-TL rely on a combination of convolutional layers, bidirectional long short-term memory (BiLSTM) layers, and a single-head attention (SHA) layer (deepTFBS backbone; **Figure 1B**). First, the deepTFBS backbone uses a convolutional layer to scan one-hot encoded input sequences, enabling the recognition of TF binding motifs. To enhance the effectiveness of the convolutional layer, and to avoid overfitting, batch normalization is applied to re-center and re-scale feature maps. Subsequently, three stacking convolutional layers are used to extract higher-level feature representations. To address issues associated with training very deep neural networks, deepTFBS uses a residual block with shortcut connection, an approach previously used in computer vision [31] and genomic signal prediction tasks [32]. The use of the residual block mitigates issues associated with vanishing gradients and performance degradation. Following the extraction of local sequence features by the convolutional layer and the residual block, a bidirectional LSTM layer is employed to capture local and global dependencies within DNA sequences. This step enables deepTFBS to extract comprehensive contextual information, further enhancing the representation of features relevant to TF binding. To detect relevant long-range dependencies and identify other features critical for TF binding, deepTFBS incorporates a SHA layer, which allows the model to focus on relevant information. Finally, a fully connected layer is utilized to make predictions by integrating features identified via convolution, BiLSMT and self-attention.

### deepTFBS-MT shows superior performance in predicting TFBSs in *Arabidopsis*

We first assessed the performance of deepTFBS-MT by training it with 359 *Arabidopsis* TF binding profiles (**Supplemental Table S1**), which were compiled from previously published studies [4, 6, 33–35]. To ensure consistent peak regions across the dataset, we employed a standardized computational pipeline (**Supplemental Figure 1**) to reprocess raw reads. Predicting TFBSs is inherently challenging since TF binding regions constitute only a minor fraction of the entire genome [4, 8]. To generate training datasets for deepTFBS-MT, we partitioned the *Arabidopsis* genome into 595,710 non-overlapping 200-bp genomic windows, extending these windows to 1,000bp. Each 1,000bp genomic window was then assigned a 359-dimensional label corresponding to the 359 TFs, based on peak regions identified by ChIP-Seq or DAP-Seq (see **METHODS**). To avoid including regions with low binding evidence that may introduce noises into the analysis, we used 295,955 genomic windows supported by at least least 5 TFs within a 1,000 bp region. These windows were split into training, validation, and testing sets for model training, hyperparameter tuning, and final evaluation, respectively (**Figure 1C**). In this context, “positive” samples represented sequences containing binding site(s) for a specific TF, while “negative” samples were sequences with binding sites associated with some of the remaining 358 TFs.

Using the hold-out testing dataset, containing 54,866 sequences, deepTFBS-MT showed an impressive performance with a median AUROC (area under receiver operating characteristics curve) of 0.964 (**Figure 2A and Supplemental Table S2**). However, considering the extreme imbalance between positive and negative samples within the training, validation, and testing datasets, the AUROC metric can potentially over-estimate model performance, which is a common limitation in imbalanced classification problems in machine learning [36]. In contrast, PRAUC (area under the precision-recall curve) does not consider true negative, therefore it is more suitable for assessing model performance when using highly imbalanced datasets where positive samples represents a minority. When deepTFBS-MT was evaluated by calculating PRAUC, using the same test dataset, we observed a median PRAUC value of 0.629 (**Figure 2B**). There were 124 TFs exhibited relatively low PRAUC values (e.g., less than 0.5), this was largely attributed to the low positive-to-negative ratio in the testing dataset (**Supplemental Figure 2**A). For example, RGA (AT2G01570), which has the lowest PRAUC of 0.0093, is represented by only 222 binding sites among a large number of negatives in the testing set (**Supplemental Table S2**). Notably, when the negative samples for RGA were downsampled to balance the dataset, the PRAUC increased significantly (**Supplemental Figure 2B**), indicating that the low PRAUC is a reflection of data imbalance rather than model performance. Similarly, for another TF, ATNAC6 (AT5G39610), with a PRAUC of 0.5031 and 1,082 binding sites, we observed a remarkably improvement in PRAUC upon downsampling negative samples (**Supplemental Figure 2C**). These findings highlight the capability of deepTFBS-MT in TF binding predicting, even when analyzing TFs with sparse binding data.

**Figure 2.**
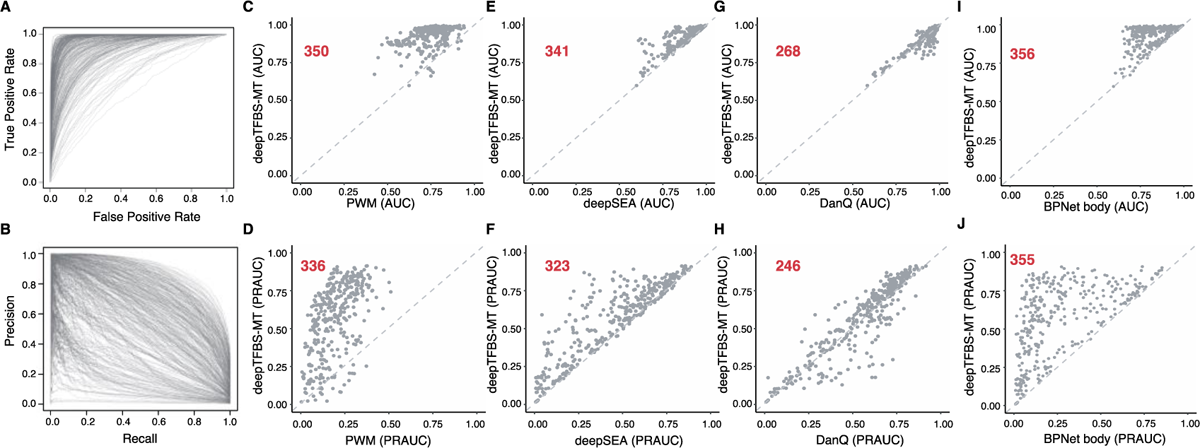
Performance evaluation of deepTFBS-MT in predicting TF binding sites in Arabidopsis. (A) Receiver operating characteristic (ROC) curves and (B) precision-recall (PR) curves for predictions across 359 TFs in *Arabidopsis*. Each curve represents one TF. (C-J) Performance comparison of deepTFBS-MT with existing methods. Scatter plots showing the comparison of area under ROC curve (AUC) between deepTFBS-MT and (C) PWM, (E) deepSEA, (G) DanQ, and (I) BPNet body. Area under PR curve (PRAUC) comparisons between deepTFBS-MT and (D) PWM, (F) deepSEA, (H) DanQ, and (J) BPNet body are also shown. Each point represents one TF, with points above the diagonal indicating superior performance by deepTFBS-MT.

Next, we compared the performance of deepTFBS-MT with the previously reported PWM-based method to demonstrate the predictive power of the new model. This approach scores genomic sequences with a matrix representing the log likelihood of each nucleotide being present in a given TF binding motif. In this analysis, we scored each DNA sequence from the *Arabidopsis* test dataset (see **METHODS** section) to ensure a fair comparison between the two approaches. By utilizing AUROC and PRAUC metrics, we found that deepTFBS-MT demonstrated superior performance for 350 out of the 359 TFs (**Figure 2C-2D**). Significantly, compared to PWM, deepTFBS-MT the median PRAUC improved from 0.189 to 0.629 (**Supplemental Table S2**). Besides PWM-based method, we also benchmarked the performance of deepTFBS against three other DL methods— DeepSEA [17], DanQ [18] and BPNet [23]. Both DeepSEA and DanQ were originally designed as classification models to predict regulatory elements (e.g., TF binding sites, histone mark, etc) in the human genome, making them directly comparable to our deepTFBS-MT after re-training on *Arabidopsis* TF-binding datasets. BPNet, which was originally designed as a regression model for base-pair resolution prediction of TF binding intensity using ChIP-nexus data. We adapted its core architecture, BPNet body, which consists of 11 stacked convolutional layers, to perform our multi-class classification task while maintaining its powerful feature extraction capability. Compared to DeepSEA, deepTFBS-MT showed a better performance in identifying the binding sites of 341 out of the 359 TFs when assessed by AUC, while it showed better performance with 323 TFs when the PRAUC measurement was used. On average, deepTFBS-MT achieved a 3.84% AUC and a 22.14% PRAUC performance gain (**Figure 2E-2F**). Similarly, when compared with DanQ, we noted that in deepTFBS-MT outperformed DanQ for 268 TFs when using AUC and 246 TFs with PRAUC metrics (**Figure 2G-2H**). When compared to the adapted BPNet architecture, deepTFBS-MT demonstrated consistently better performance across both AUC and PRAUC metrics (**Figure 2I-2J**), indicating that our hybrid architecture of CNN, BiLSTM, and attention layers is more effective for predicting TF binding patterns.

To investigate the contributions of individual components to the model’s performance, we conducted a systematic ablation study by training six model variants: (1) CNN only, (2) LSTM only, (3) single-head attention (SHA) only, (4) CNN+LSTM, (5) CNN+SHA, and (6) LSTM+SHA. As shown in **Figure 3**, the full architecture of CNN+LSTM+SHA, which underpins deepTFBS, achieved the highest median PRAUC, highlighting the synergistic effect of these components. In contrast, the LSTM-only architecture showed the poorest performance, underscoring the importance of CNN for extracting local patterns. While the improvement from CNN+LSTM (PRAUC: 0.602) to CNN+LSTM+SHA (PRAUC: 0.629) was modest, it demonstrates that SHA adds value, albeit incrementally. These results demonstrate the efficacy of the deepTFBS-MT architecture in accurately predicting TFBSs in Arabidopsis, providing a robust framework for transcription factor binding prediction.

**Figure 3.**
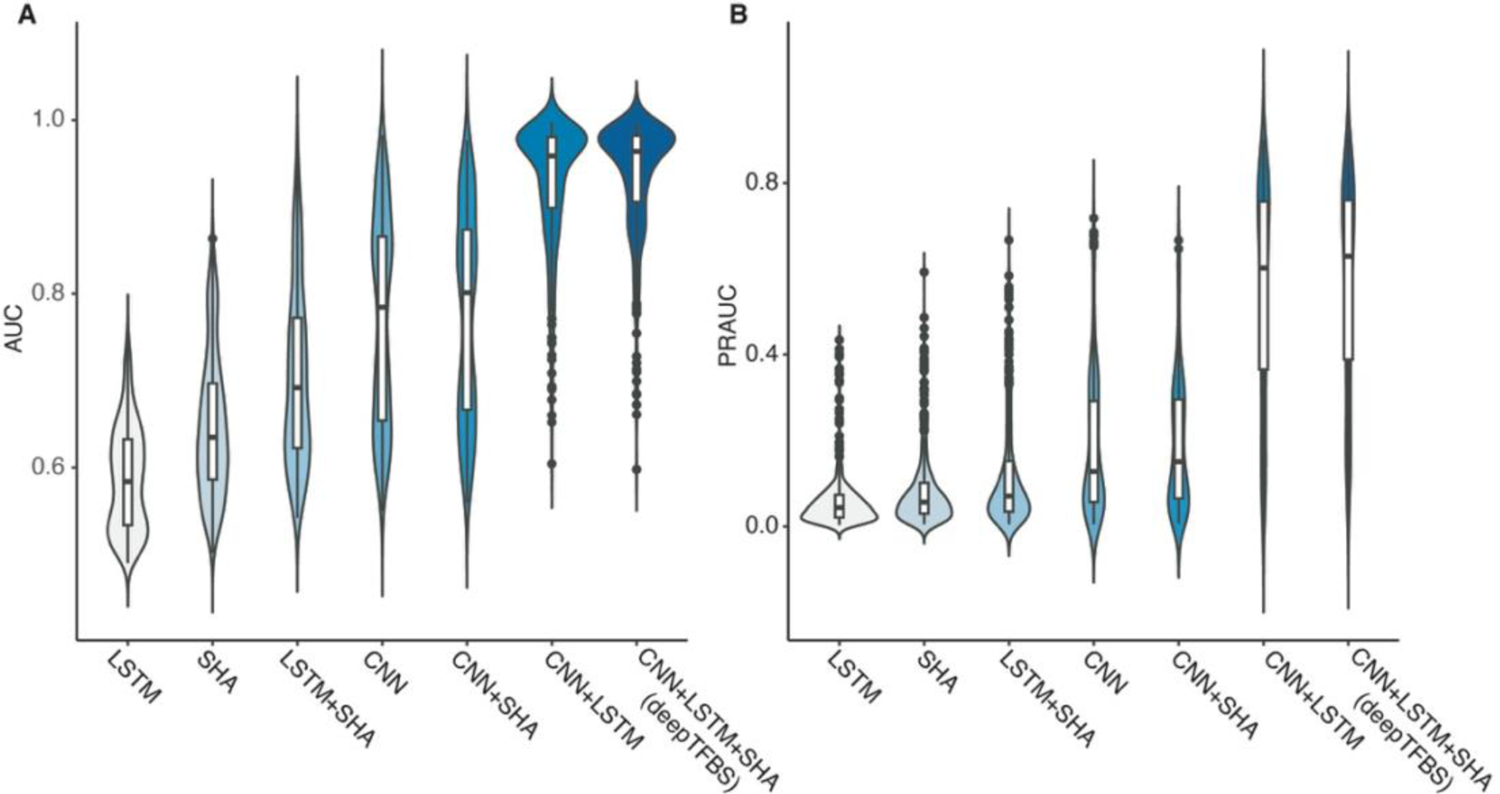
Ablation study demonstrating the contribution of different architectural components in deepTFBS-MT. (A) Area under ROC curve (AUC) and (B) area under precision-recall curve (PRAUC) for different model variants. Seven architectures were evaluated: LSTM only, single-head attention (SHA) only, LSTM+SHA, CNN only, CNN+SHA, CNN+LSTM, and the complete deepTFBS architecture (CNN+LSTM+SHA). Violin plots show the performance distribution across all TFs, with box plots indicating the median and quartiles. The complete architecture (CNN+LSTM+SHA) achieved the highest median performance, while LSTM-only showed the lowest..

### deepTFBS-MT has interpretation ability by identifying TF binding preferences

To better understand the factors influencing deepTFBS-MT’s performance, we applied an Integrated Gradients (IG) [37, 38], a gradient-based attribution method, to visualize the importance of individual features identified by the model (see **METHODS** section). IG computes the average gradient of the model’s output as input features change, quantifying the importance of each nucleotide. A high absolute gradient for a feature indicates its greater significance in the prediction process. Using this method to sequences in the test dataset, we identified putative motifs for the top ten TFs predicted by deepTFBS-MT. Notably, seven of these ten motifs were also recorded in the Plant Cistrome Database [4]. Based on PRAUC metrics, the highest ranked TF was CBF4, known for its role in drought stress [39]. The consensus motif identified by deepTFBS-MT for CBF4 was [TC]GTCGG, which closely matches the motif recorded in the Plant Cistrome Database ([ATC][TC]GTCGG[CT]) (**Figure 4A**). Similarly, AT3G16280 and CBF2, both members of the AP2-EREBP family, displayed nearly identical motifs, suggesting shared TF binding preferences among these TFs. For the remaining TFs—EPR1, GBF3, ATHB25, and AP2—the motifs identified by deepTFBS-MT closely aligned with those in the Plant Cistrome Database (**Figure 4A**). These results indicate that deepTFBS-MT has the capability of learning the binding preferences of different TFs.

**Figure 4.**
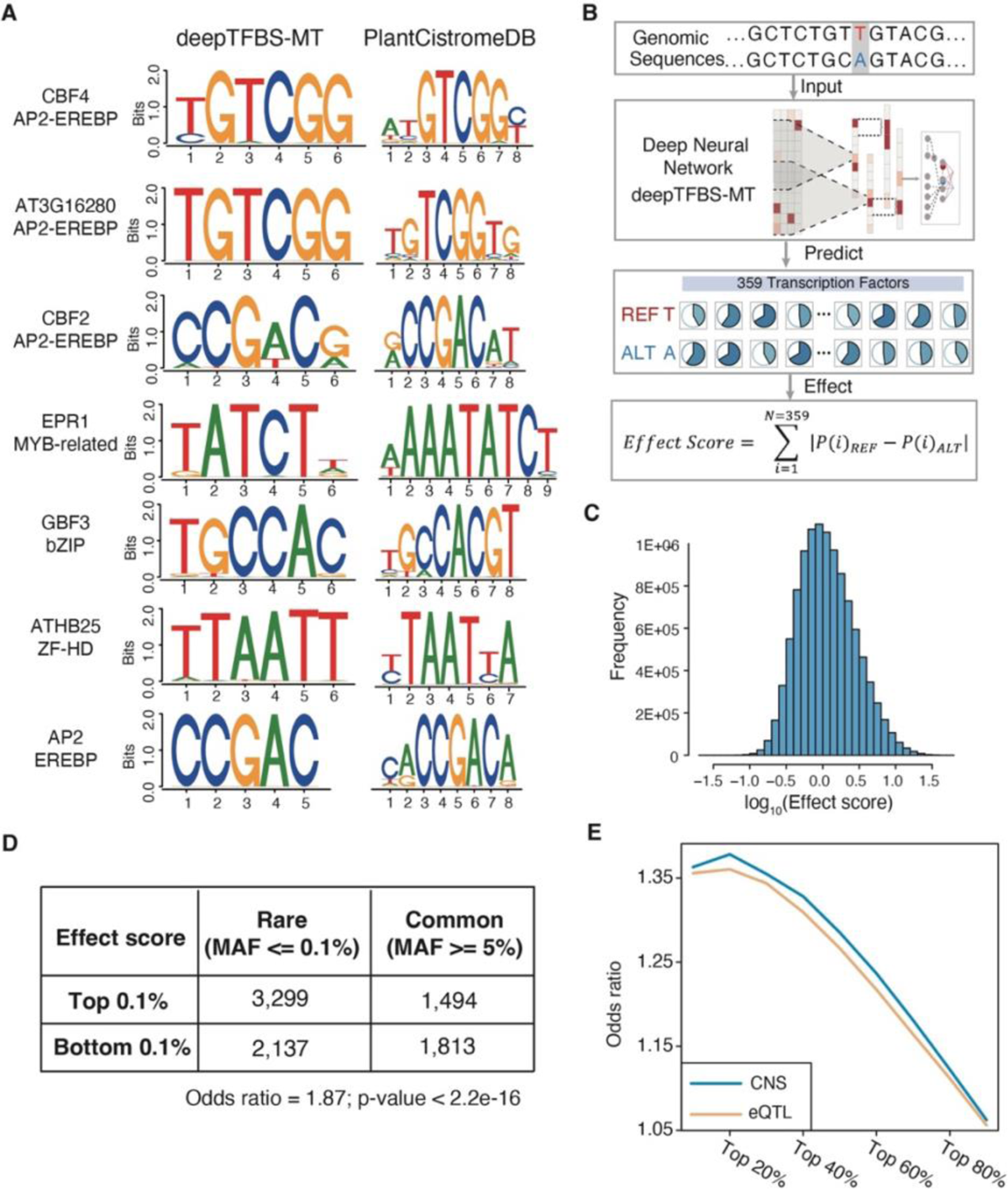
Interpretation of deepTFBS-MT predictions and analysis of regulatory variants. (A) Comparison of binding motifs identified by deepTFBS-MT (left) and those documented in PlantCistromeDB (right) for the top-performing TFs. Sequence logos show the position-specific nucleotide preferences, with height indicating information content. (B) Schematic illustration of the regulatory variant effect prediction pipeline. The process includes input sequence pairs (reference and alternate alleles), prediction of binding probabilities for 359 TFs using deepTFBS-MT, and calculation of an aggregate effect score based on differential binding predictions. (C) Distribution of effect scores across all analyzed SNPs from the 1001 Genomes Project (n=10,706,842), shown on a log10 scale. (D) Enrichment analysis of rare (MAF ≤ 0.1%) versus common (MAF ≥ 5%) variants among SNPs with high (top 0.1%) and low (bottom 0.1%) effect scores. The odds ratio of 1.87 indicates significant enrichment of rare variants among high-impact SNPs was determined by Fisher’s exact test. (E) Odds ratio plots showing the enrichment of predicted high-impact variants in conserved non-coding sequences (CNS, blue) and expression quantitative trait loci (eQTL, orange) across different effect score thresholds.

### deepTFBS-MT predicts putative functional regulatory variants

Given the efficacy of deepTFBS in understanding TF binding grammar, we next investigated whether it could be applied to identify candidate functional regulatory variants, which are known to play a crucial role in complex phenotypes [40, 41]. To assess the regulatory impact of individual SNPs, we analyzed sequence pairs comprising both reference and alternate alleles, each centered within their flanking regions. These sequences were input into the deepTFBS-MT model to predict the binding probabilities of 359 TFs for each allele. The regulatory impact of each SNP was then quantified by computing the absolute differences in predicted binding scores between reference and alternate alleles. We defined an aggregate ‘effect score’ for each SNP by summing these differential binding probabilities across all 359 TFs (**Figure 4B**).

We applied this analytical approach to 10,706,842 SNPs from naturally occurring accessions in the 1001 Genomes Project [42]. As expected, the majority of these SNPs had very low effect scores (**Figure 4C**). However, when focusing on the top 10% of SNPs with the highest effect scores, we found that over 66% were located in either promoter (upstream 1000bp) or distal intergenic regions, suggesting a potential role in gene regulation (**Supplemental Figure S3A**). To further validate deepTFBS-MT’s ablity to identify putative functional regulatory variants, we assessed the enrichment of rare alleles (minor allele frequency [MAF] < 0.001) in highly impactful regulatory variants (top 0.1% of SNPs with highest effect scores) compared to those with low impact (bottom 0.1%). Since functional variants tend to have lower frequencies in populations due to selective constraints [43], we hypothesized an enrichment of rare alleles in highly impactful variants. Consistent with this hypothesis, we observed an enrichment of rare alleles with an odds ratio of 1.87 (**Figure 4D**; *p*-value < 2.2e-16 by Fisher’s exact test). Additionally, we observed that as effect scores increased, the average MAF decreased (**Supplemental Figure S3B**), indicating that highly impactful regulatory variants tend to be rarer in the population. Beyond allele frequency, we assessed the enrichment of these variants in conserved non-coding sequences (CNS) [44]. As expected, highly impactful variants showed higher odds ratios for enrichment in CNS, further supporting their potential functional relevance (**Figure 3E**). These findings demonstrate deepTFBS-MT’s ability to prioritize regulatory variants likely to play key roles in gene regulation.

When examining the highly impactful regulatory motifs, we noticed two interesting examples. The first is the SNP, chr2:6321143 (T/A), located upstream of the *PR1* gene, which is associated with leaf yellowing disease [45]. Population transcriptome data from the 1001 Genomes Project has demonstrated that this SNP affects the expression of the *PR1* gene (**Supplemental Figure 3C**). According to deepTFBS-MT predictions, the “A” allele reduced the binding affinity of multiple WRKY family TFs, including WRKY22 (AT4G01250), WRKY27 (AT5G52830), WRKY8 (AT5G46350), and WRKY15 (AT2G23320) (**Supplemental Figure 3D**). This supports previous findings that WRKY TFs interact with the *PR1* gene promoter, synergistically stimulating *PR1* expression in response to salicylic acid [46]. The second example is SNP chr3: 3059896 (A/C), associated with flowering time. According to deepTFBS-MT predictions (**Supplemental Figure 3D**), this SNP disrupts the binding motif of ATSPL9, a TF known to affect flowering in *Arabidopsis*. Taken together, these examples demonstrate how deepTFBS-MT predictions can be integrated with GWAS results to offer mechanistic insights into how specific SNPs influence gene expression and phenotypic traits, aiding the identification of causal mutations associated with complex traits.

### deepTFBS-TL shows enhanced TFBS prediction through transfer learning both within and across plant species

Although deepTFBS-MT showed impressive performance in predicting TFBSs in *Arabidopsis*, it relied heavily on large TF binding datasets. Such extensive datasets are often unavailable for other plant species, especially crops. To address this limitation, we developed deepTFBS-TL (**Figure 1A**), a transfer learning approach that uses sequence patterns learned by deepTFBS-MT from large scale TF binding profiles. By employing transfer learning, deepTFBS-TL initializes with the pre-trained weights from deepTFBS-MT, narrowing the parameter search space. As a result, deepTFBS-TL is expected to perform well even with fewer training observations. deepTFBS-TL is designed as a binary classifier. It uses the same positive samples as deepTFBS-MT and generates negative samples by selecting 10 random sets, each matching the number of positives. For each TF, we trained 10 deepTFBS-TL models using positive samples and the corresponding negative sets. To have a fair comparison to deepTFBS-MT, we evaluated deepTFBS-TL on the same imbalanced test dataset. deepTFBS-TL improved performance in 347 out of 359 TFs, achieving an average PRAUC improvement of 50.5% compared to deepTFBS-MT (**Figure 5A and Supplemental Table S4**).

**Figure 5.**
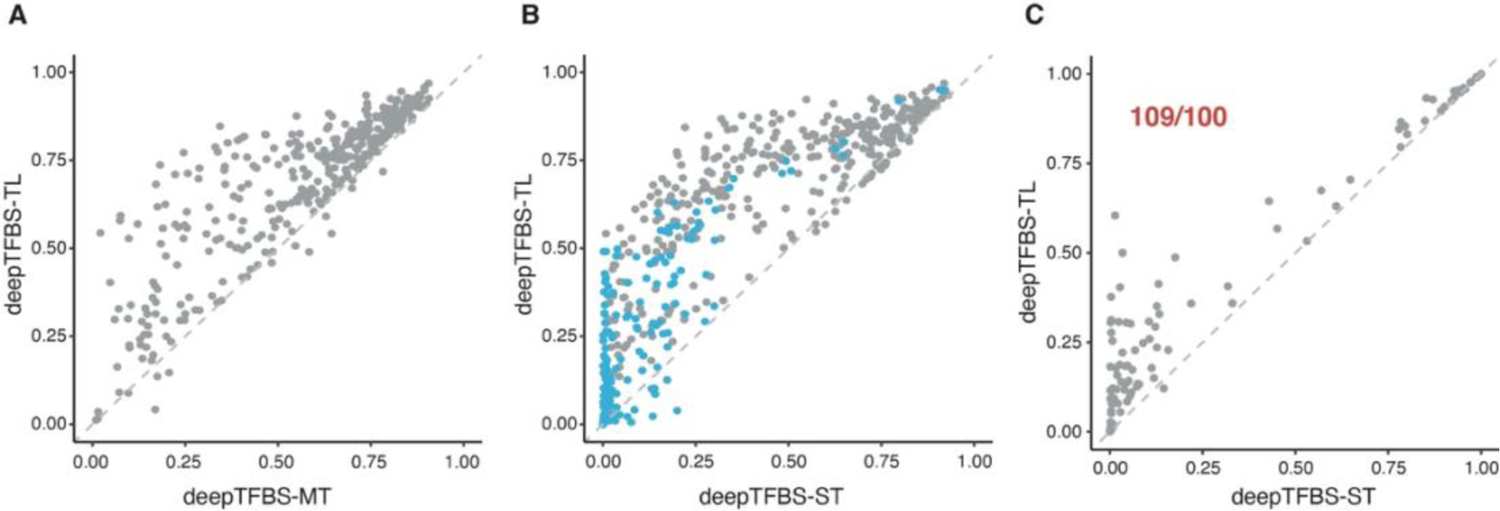
Performance comparison of transfer learning strategies in deepTFBS. (A) Scatter plot comparing PRAUC values between deepTFBS-MT and deepTFBS-TL for 359 Arabidopsis TFs. Points above the diagonal line indicate improved performance with transfer learning. (B) Performance comparison between deepTFBS-ST (trained from scratch) and deepTFBS-TL (with transfer learning) for Arabidopsis TFs. Blue points highlight TFs with fewer than 1,000 binding sites, demonstrating the particular advantage of transfer learning for TFs with limited training data. (C) Cross-species prediction performance comparison between deepTFBS-ST and deepTFBS-TL for 110 wheat TFs. deepTFBS-TL outperformed deepTFBS-ST in 109 out of 110 cases (indicated in red), demonstrating the effectiveness of transfer learning in cross-species applications.

To quantify the impact of transfer learning, we compared deepTFBS-TL, which leverages pre-trained weights from deepTFBS-MT, to deepTFBS-ST, a binary classifier trained from scratch without transfer learning. Testing on the same datasets showed transfer learning led to superior binding site prediction in 339 out of 359 TFs, resulting in an average PRAUC improvement of 160% (**Figure 5B**). We also examined how deepTFBS-TL performs when dealing with TFs that have fewer peak regions. Specifically, we created a reduced training dataset by removing 153 TFs with fewer than 1,000 peak regions, as detailed in the **METHODS** section. We then trained both models— deepTFBS-TL with transfer learning and deepTFBS-ST without it—on these TFs. The results demonstrated that deepTFBS-TL significantly improved binding site identification for 135 out of 153 TFs, achieving an average PRAUC improvement of 16.9% (**Figure 5B**). However, when analyzing the performance across different TFs revealed a slight negative correlation (*R* = −0.13 and *P*-value = 0.0024) between PRAUC improvement and the number of binding sites used in training (**Supplemental Figure 4A**), emphasizing that transfer learning is especially beneficial for models trained on smaller sample sizes.

Next, we assessed the cross-species predictive capabilities of deepTFBS-TL using curated binding profiles for 110 TFs in wheat (**METHODS**). Utilizing the pre-trained deepTFBS-MT, we trained 110 deepTFBS-TL and deepTFBS-ST binary classifiers specifically for wheat. When evaluated on the same *Arabidopsis* test set (**Figure 1C**), deepTFBS-TL outperformed deepTFBS-ST in 109 out of 110 TFBS predictions (**Figure 5C; Supplemental Table S4**). We also observed a trend where deepTFBS-TL showed slightly greater improvement for TFs with fewer binding sites (**Supplemental Figure 4B**), suggesting a potential benefit of transfer learning in cases with limited training data, even across species. Overall, these results highlight the effectiveness of deepTFBS-TL, demonstrating that transfer learning significantly enhances TFBS prediction performance, especially in scenarios with limited training data and across different plant species.

### Application of deepTFBS to study the *WUS* regulatory network in *Arabidopsis*, maize, and wheat

Given the robust performance of deepTFBS in both intra- and inter-species predications, as a test case, we applied this framework to explore *WUSCHEL* (*WUS)* regulatory network. WUS is a TF crucial for stem cell maintenance and floral development [47]. We generated *WUS* DAP-Seq data in *Arabidopsis* and maize, analyzing the results with a consistent data processing pipeline (**Supplemental Figure 1**). In these experiments we obtained two biological replicates which generated consistent peaks, and the analysis of the genomic distribution of TFBSs revealed an enrichment of binding sites in intergenic regions (**Supplemental Figure 5** and **6**). In our deepTFBS-TL tests, the *WUS*-target classifiers derived from *Arabidopsis* and maize, showed a PRAUC of 0.651 and 0.671, respectively (**Figure 6A**).

**Figure 6.**
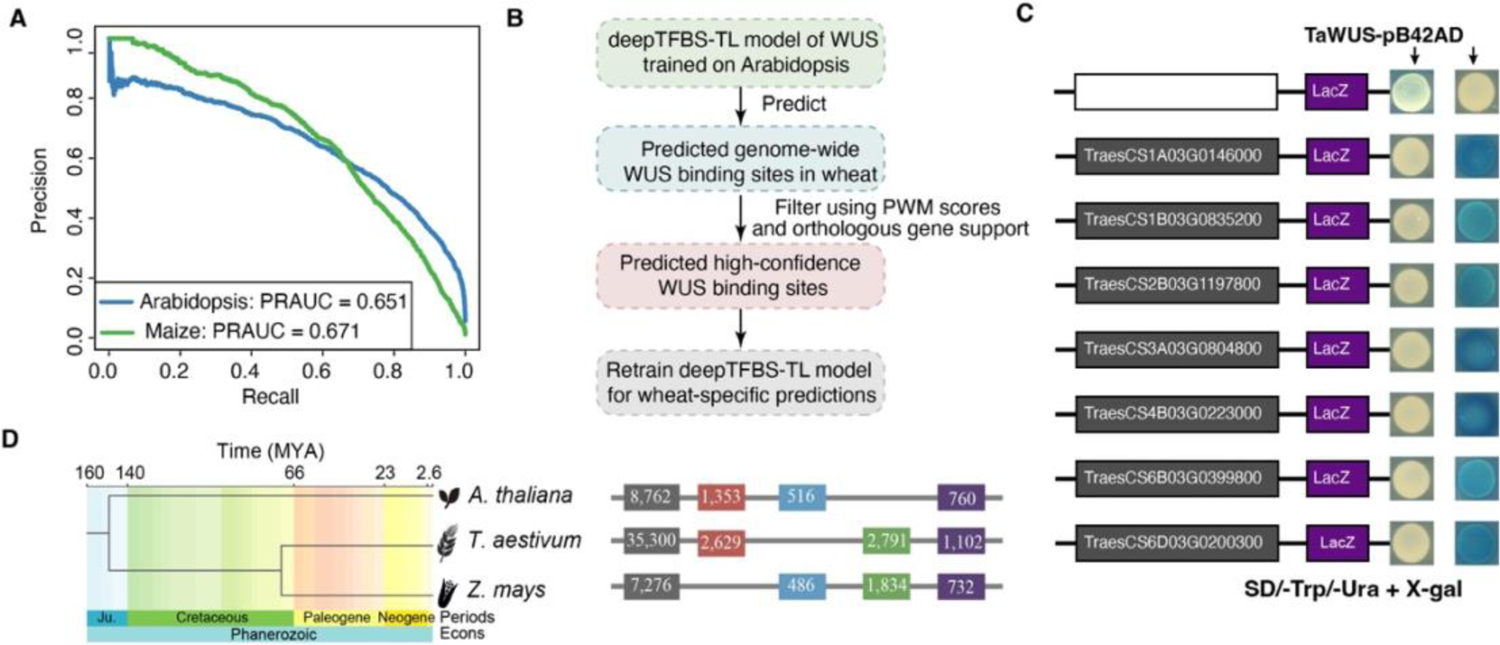
Application of deepTFBS to study WUS regulatory networks across plant species. (A) Precision-recall curves showing the performance of deepTFBS-TL in predicting WUS binding sites in Arabidopsis (blue, PRAUC = 0.651) and maize (green, PRAUC = 0.671). (B) Workflow for predicting WUS binding sites in wheat. The process begins with a deepTFBS-TL model trained on Arabidopsis data, followed by genome-wide prediction in wheat, filtering using PWM scores and orthologous support, and retraining for wheat-specific predictions. (C) Experimental validation of predicted WUS binding sites using yeast one-hybrid (Y1H) assays. Seven representative positive interactions are shown, with blue colonies indicating positive binding events. The empty vector (pB42AD) serves as a negative control. (D) Evolutionary conservation analysis of WUS binding sites. Left: phylogenetic tree showing divergence times (MYA) between Arabidopsis, wheat, and maize. Right: horizontal bars showing the number of predicted WUS binding sites in each species and their conservation patterns (colored segments represent different conservation categories).

To generate genome-wide TFBS data in scenarios where DAP-Seq data for a specific TF is unavailable in a plant species, we developed a process summarized in **Figure 6B**. Using wheat as an example, we initially applied deepTFBS-TL, trained on *Arabidopsis* data, to predit TFBS in the wheat genome. We then employed FIMO [48] to scan and analyze the identified genomic regions to determine if the existence of the candidate regions was supported by *Arabidopsis* PWM data. Furthermore, orthologous targets from *Arabidopsis* were used to substantiate the validity of the predictions. The sites predicted and supported by either PWM or orthologous target sequences were then used as “positives” for training deepTFBS-TL classifiers. To verify cross-species prediction validity, we randomly selected 14 high-confidence targets identified in wheat for testing with yeast one-hybrid (Y1H) assays, confirming the binding of seven out of 14 predicted targets experimentally, thereby demonstrating our prediction method’s reliability (**Figure 6C**). In contrast, when we randomly selected 16 predicted targets based solely on PWM, only two were identified as positives (**Supplemental Figure 7 and Supplemental Table S5**).

To assess the evolutionary conservation of *WUS* targets across *Arabidopsis*, maize and wheat, we developed species-specific deepTFBS-TL models to predict genome-wide DNA binding targets. Comparative analyses of the predicted TFBSs showed a moderate level of conservation, with 760 targets in the *Arabidopsis* genome conserved across all three species (**Figure 6D and Supplemental Table S6-S8**). Notably, several experimentally validated WUS targets, such as TPR1 and TPR2—essential for embryonic patterning and auxin response [47, 49]—and ATAVP1, involved in various developmental processes [47, 50], were among these conserved targets. This finding validates our computational approach and undersores the biological significance of the conserved binding sites. Gene Ontology analysis of the 760 conserved targets highlighted their roles in diverse biological processes, including ion transport and metabolism, pollen development and cellular pH regulation (**Supplemental Figure 8** and **Supplemental Table S6**), reflecting the multifaceted functions of WUS in plant development.

### deepTFBS is available as an open source framework and a web server

To facilitate its broad application, and provide enhanced functionality for plant biologists, we made the source code for deepTFBS publicly available on GitHub (https://github.com/cma2015/deepTFBS). In addition, to mitigate potential inconsistencies arising from dependencies in a DL framework, a Docker image was also created and made publicly accessible (https://hub.docker.com/r/malab/deeptfbs). Furthermore, to accommodate experimental biologists lacking programming expertise, a web-based interface was also developed to facilitate the prediction of TFBSs in any given DNA sequence. The web server comprises of two primary components: a database and a predictor option. Within the database users can query and retrieve predicted binding sites for 512 *Arabidopsis* and 110 wheat TFs (**Supplemental Figure 9**). On the other hand, the TFBS prediction feature accepts any DNA sequence as input and generates a map of predicted TFBSs via an interactive graphical interface that highlights positive predicted binding sites. Each online prediction request is assigned a unique job identifier, allowing users to monitor the status of their prediction requests. However, for large-scale prediction requirements, we highly recommend the utilization of the deepTFBS Docker image to ensure scalability (https://hub.docker.com/r/malab/deeptfbs).

## DISCUSSION

The accurate identification of TF binding sites is essential for elucidating intricate transcriptional regulatory network features. We introduce deepTFBS, a framework that improves upon traditional PWM-based methods by capturing long-range regulatory interactions, while addressing the limitations of current DL approaches through transfer learning. The framework’s core architecture combines stacked CNN layers, a BiLSTM layer, and a self-attention layer to process DNA sequences (**Figure 1**). Compared to existing DL methods such as DeepSEA and DanQ, the deepTFBS framework first leverages multi-task learning (deepTFBS-MT) to simultaneously predict binding specificities of multiple TFs in Arabidopsis, optimally utilizing large datasets and capturing shared information among TFs. To address scenarios with limited data availability, we then developed deepTFBS-TL. By employing transfer learning strategies, this approach effectively tackles the sample size constraints and enables accurate TFBS predictions across different plant species, making the framework versatile and practical in broader applications.

deepTFBS shows potential to identify putative functional regulatory variants, which may contribute to complex phenotypes. By analyzing sequence pairs of reference and alternate alleles for over 10 million SNPs from the 1001 Genomes Project, we found that deepTFBS-MT can quantify the regulatory impact of genetic variants through an aggregate effect score across 359 TFs. Several observations suggest the biological relevance of these predictions: highly impactful variants are enriched in promoter and distal intergenic regions, show higher frequencies of rare alleles (OR = 1.87), and are preferentially located in conserved non-coding sequences. Examination of specific cases, such as the SNPs near the *PR1* gene and those associated with flowering time, suggests how deepTFBS-MT predictions could help understand the relationship between genetic variation and phenotypic traits. The integration of these predictions with GWAS results may provide insights into potential regulatory mechanisms underlying complex traits.

The application of deepTFBS to investigate *WUS* regulatory networks across *Arabidopsis*, maize and wheat undersores its effectiveness in comparative regulatory studies. By utilizing DAP-seq data from *Arabidopsis* and maize, we achieved respectable prediction accuracies, with PRAUC values of 0.651 and 0.671, respectively. Our integrated approach for wheat, which combines deepTFBS-TL predictions with PWM and orthologous information, yielded encouraging results in Y1H validation experiments (**Figure 6**). The discovery of 760 conserved WUS targets across these three species, including previously validated targets such as TPR1, TPR2, and ATAVP1, highlights deepTFBS’s potential to reveal evolutionarily conserved regulatory relationships. This study suggests that deepTFBS could serve as a valuable tool for exploring transcriptional regulation across plant species, especially in contexts where experimental binding data is sparse.

Overall, deepTFBS offers a robust and adaptable framework for exploring gene regulation in plants, particularly through its ability to integrate large-scale TF binding profiles and provide accurate predictions across species. The framework should especially valuable for studying transcriptional regulation in non-model plants where binding data is limited. To facilitate the application of deepTFBS, we provide the source codes, a web server, a Docker image, and comprehensive user documentation to the scientific community (https://github.com/cma2015/deepTFBS).

## METHODS

### *Arabidopsis* and wheat TF binding data

*Arabidopsis* TF binding data, including ChIP-Seq and DAP-Seq results, were collated from five different sources: 1) The PlantTFDB database [33] that provides ChIP-Seq datasets for 9 TFs; 2) PlantCistromeDB [4] containing DAP-Seq datasets for 327 TFs, providing a valuable resource for plant cistrome data; 3) Six TFs and their corresponding binding data were derived from PCSD [34]; 4) A study describing ChIP-Seq datasets for 21 TFs in exploring the role of ABA in plant growth, development, and stress responses [6]; and 5) DAP-Seq datasets for 3 TFs aiming to characterize the TF binding landscape in *Arabidopsis* [51]. For wheat, DAP-Seq datasets for 110 TFs were obtained from a recent study by [9], providing valuable insights into the regulatory networks of this economically important crop species.

### Illumina library preparation and *WUS* DAP-Seq in *Arabidopsis* and Maize

All the plant seed were surface-sterilized in a 10% sodium hypochlorite (NaClO) solution. Arabidopsis seeds were then germinated on half-strength Murashige-Skoog (1/2 MS) medium supplemented with 10% sucrose and 1% agar (w/v). Plants were cultivated in a controlled growth chamber with a 16-hour light/8-hour dark photoperiod, maintaining a temperature of 22℃ during the light period with an illumination intensity of 80–90 µmol m−2 s−1, and 18℃ in the dark phase. Wheat (Chinese spring) seeds,were germinated and grown on half-strength Hoagland’s liquid medium under long-day conditions (16 h light and 8 h dark) at 22°C with 50% relative humidity and a photon flux density of 300μmol m−2s−1. The maize (B73) seeds were germinated in a culture room set with a 12-h photoperiod, maintaining a photon flux density of 8000 LUX and a constant temperature of 28 ± 1°C. Genomic DNA was extracted from whole seedlings of *Arabidopsis* and maize and 1 μg of this DNA was uniformly fragmented to an average size of 200 bp using the Bioruptor Plus sonicator (Diagenode). Following fragmentation, DNA ends underwent an end-repair process, and an A-tail was added employing the NEXTflex Rapid DNA-Seq Kit (Revvity). Next, adapters were ligated to the A-tailed DNA fragments, preparing them as libraries for downstream DAP-Seq analysis. The DAP-Seq experiments were conducted in two independent replicates. The raw DAP-Seq data have been deposited in CNCB (https://www.cncb.ac.cn) under project PRJCA033158.

### Bioinformatics processing of DAP-Seq and ChIP-Seq datasets

Raw *Arabidopsis* DAP-Seq or ChIP-Seq datasets were downloaded from the NCBI’s SRA Database (https://www.ncbi.nlm.nih.gov/sra) and their quality control was performed using FastQC (Version 0.11.5). Low-quality reads and adaptor sequences were trimmed using fastp (v0.19.5) [52]. Bowtie (v1.1.2) [53] was used to align clean sequencing reads to the TAIR10 *Arabidopsis* reference genome, using default parameters. Enriched regions were identified using the MACS2 software (v2.1.1) [54] with default parameters, except the “-g” parameter, which was set to the genome size (1.25e8 for *Arabidopsis*). For custom *WUS* DAP-Seq datasets, the same pipeline was followed. Genome-wide peak visualization and annotation were performed using the R package “ChIPseeker” [55].

### Sample generation and feature encoding

To prepare the input for the training of deepTFBS, the *Arabidopsis* (TAIR10) and wheat (IWGSC v1.0) genomes were divided into non-overlapping 200-bp bins. Each of these bins was assigned to a 359-dimensional label reflecting its overlap with known TF binding peak regions. In this process, a label of “1” was assigned if the overlapping region exceeded 100 bp, otherwise, a “0” label was given. Similar procedure was followed for the wheat genome. Subsequently, for each 200-bp sequence in a bin, 400 bp uptream and downstream flanking sequences were extracted from the *Arabidopsis* and wheat genomes, resulting in a total sequence length of 1,000bp and one-hot encoding was used to convert the DNA sequences into a matrix with 1,000 rows and four columns, with each column representing one of the four nucleotides as follows:

*A* = [1, 0, 0, 0]

*C* = [0, 1, 0, 0]

*G* = [0, 0, 1, 0]

*T* = [0, 0, 0, 1]

*N* = [0, 0, 0, 0]

When evaluating the performance of deepTFBS, to avoid overfitting the *Arabidopsis* and wheat datasets were divided into three subsets: a “training,” a “validation,” and a “testing” set. The training datasets were used to train deepTFBS, the validation datasets for tuning hyperparameters, while the test datasets were used to evaluate model performance.

### The deepTFBS architecture and training

#### The backbone of deepTFBS

The backbone of deepTFBS is a hybrid network structure that combines a residual neural network with BiLSTM. The initial layer is a convolutional layer that applies one-dimensional convolution operations with a specified number of kernels (weight matrices). The outputs of these operations are then processed through the rectified linear activation function (ReLU), setting values below 0 to 0. The second part of deepTFBS consist of a residual neural network, utilized to avoid gradient vanishing. In the following step, deepTFBS employs BiLSTM to scan for long-range sequence interactions, thereby capturing more comprehensive sequence features. Then self-attention layers were used to directly focus on critical features across the entire sequence. The architecture concludes with two fully connected layers. The first one of these incorporates Dropout to mitigate overfitting by randomly omitting some features, while the number of neurons in the final output layer corresponds to the number of TFs being tested.

#### deepTFBS-MT pretraining

deepTFBS-MT was pretrained on 359 Arabidopsis TF binding datasets. The output layer contains 359 neurons corresponding to Arabidopsis TFs. The model was optimized using Adam optimizer with a batch size of 128, binary cross-entropy loss function, and early stopping (patience=10, maximum 200 epochs).

#### Fine-tuning deepTFBS-TL with transfer learning

After pre-training deepTFBS-MT, deepTFBS-TL fine-tunes the model for each specific TF using a balanced dataset approach. For each TF, we maintain all positive binding sites while randomly sampling an equal number of negative sequences to create 10 different balanced datasets. These datasets share the same positive examples but have different negative sets, enabling the training of 10 independent models per TF. deepTFBS-TL initializes each model using the pretrained weights from deepTFBS-MT, replacing only the final output layer of deepTFBS-MT with a single neuron for binary classification. This ensemble approach with multiple negative sets helps capture binding site variability while reducing bias from negative sequence selection. For comparison, deepTFBS-ST follows the identical training procedure but initializes all weights randomly instead of using pretrained weights. All models were implemented using TensorFlow (v2.6) on servers with dual NVIDIA GeForce 3090Ti GPUs (24G memory; 10,752 CUDA cores).

### Model evaluation

#### ROC

The receiver operating characteristic (ROC) curve is a widely used tool for evaluating the performance of binary classification models. The ROC curve is generated by plotting the true positive rate on the *y*-axis against the false positive rate on the *x*-axis at varied thresholds. The AUC value is then used to quantitatively assess the prediction accuracy of the binary model. The AUC value ranges from 0 to 1, with a higher AUC value indicating better prediction accuracy.

#### PRAUC

The PRAUC metric represents the area under the precision-recall curve. It is constructed by plotting precision (*y*-axis) against recall (*x*-axis) at varied thresholds. Similar to the ROC curve, the area under the precision-recall curve (PRAUC value) provides a quantitative measure of the accuracy of a binary model. The PRAUC value also ranges from 0 to 1, with a higher values denoting superior prediction accuracy.

#### Comparison with PWM, DeepSEA, and DanQ

In order to determine the performance of deepTFBS, a traditional PWM-based method, two DL methods, and a machine-learning based method were used for comparison purposes.

### PWM-based method

Given sequence S with the length *L*, and a PWM matrix *X* _*i, j*_, where *i* = *A*, *C*, *G*, *T*, *j* represents the position of corresponding nucleotide, *j* = 1,2, …, *n*, PWM can be represented as follows:

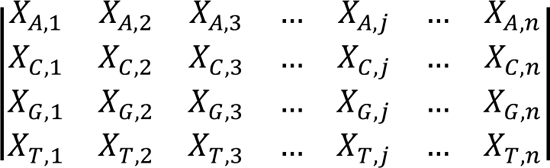

Each sequence can be split into *m k*-mers and then *k*-mers can be scored as

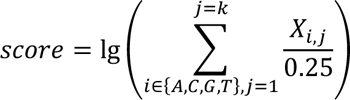

#### DeepSEA

DeepSEA [17] is a multi-label classification model based on a CNN, originally utilized for predicting functional noncoding DNA variants in humans. However, DeepSEA could not be directly utilized to predict TFBSs in plant species. Therefore, based on the network structure and hyperparameters outlined in the original publication, DeepSEA was re-trained using the *Arabidopsis* TFBS datasets.

#### DanQ

Just as DeepSEA, DanQ [18] is a multi-label classification model that combines a CNN with LSTM. It was initially trained on the same data as DeepSEA. Utilizing Keras, we re-trained and then used DanQ with the same training, validation, and testing datasets as those used for deepTFBS, ensuring the consistency of performance comparisons.

### Model interpretation by integrated gradients and consensus motif identification

Gradient-based attribution methods, for example IG, are widely used to explore the contribution of input features to the output of deep neural networks [37]. IG calculates the gradients of the output neuron in response to input features, measuring how the output changes with small perturbations in the input. These methods assign an attribution value to each input feature, representing its contribution to the output. Applying this method to deepTFBS, the target neurons represent individual TFs, and the IG method is employed to compute the average gradient of the output, effectively mitigating the saturation problem that arises when calculating gradients at the input layer. By calculating the attribution value for each position in the input sequence, and normalizing the results, it becomes possible to visualize the influence of each nucleotide on the prediction. Higher attribution values indicate a greater importance of a nucleotide at the corresponding position within the evaluated TFBS. The implementation of IG was via deepExplain (https://github.com/marcoancona/DeepExplain).

For each TF, attribution values were calculated for sequences in the test set when predicted scores exceeded 0.9, using the IG method. These attribution values were used to quantify the importance of each position within the input sequence on the prediction performance of the model. For a sequence of *L*, the top k motifs were searched based on the magnitude of attribution values, generating a comprehensive set of candidate motifs for each TF. To determine the consensus motif, the attribution values were averaged within each window, and the highest-scoring motif was selected while its neighbors were removed to avoid redundancy. The UMAP algorithm was used to embed the obtained motifs into a lower-dimensional space, capturing the underlying structure and similarity between the motifs. The DBSCAN algorithm was then applied to cluster the embedded motifs to generate the consensus motif.

### Yeast one-hybrid assays

To conduct the yeast one-hybrid assays, the coding sequences of the *TaWUS* TFs was cloned individually into the pB42AD expression vector. Predicted gene promoter sequences potentially binding to *AtWUS* and *TaWUS* were inserted into separate pLacZi vectors. The recombinant pB42AD and pLacZi vectors containing the promoter sequences were co-transformed into the yeast strain EGY48A, using empty pB42AD and pLacZi vectors as negative controls. Individual clones were transferred to SD/−Trp/−Ura medium containing X-gal and cultivated at 28 °C for 3 d after selection on the SD/−Trp/−Ura plates.

## AUTHOR CONTRIBUTIONS

C.M. conceived the project. J.Z implemented the algorithm, J.Z., C.Z., M.S. and P.D. collected all the datasets and performed all the analysis. X.Y. and C.T. performed validation experiments. J.Z., C.M., Y.Z. and Z.L. interpreted data. J.Z., C.M. and Y.Z. wrote the article. All authors have read and approved the final manuscript.

## ACKNOWLEDGEMENTS

We thank the High-Performance Computing (HPC) of Northwest A&F University for providing computing resources, prof. Jifeng Ning in College of Information Engineering, Northwest A&F University for the discussions on the model design and members of the C. Ma Labs for their constructive feedback on this work.

## Supplemental Information

**Supplemental Figure 1.** Standardized computational pipeline for processing DAP-Seq and ChIP-Seq data.

**Supplemental Figure 2.** Impact of class imbalance on deepTFBS-MT performance. (A) Relationship between model performance (PRAUC) and proportion of positive samples in the test dataset for 359 Arabidopsis TFs. Each point represents one TF, showing lower PRAUC values generally correspond to TFs with fewer positive samples. (B) Precision-recall curves for RGA (AT2G01570) at different negative-to-positive (N/P) ratios. The model’s performance decreases significantly as class imbalance increases, with PRAUC dropping from 0.677 (N/P=1) to 0.023 (N/P=100). Dotted lines represent individual runs; solid lines show the average. (C) Precision-recall curves for ATNAC6 (AT5G39610) across varying N/P ratios, demonstrating similar trend of performance decline with increasing class imbalance. PRAUC decreases from 0.955 (N/P=1) to 0.542 (N/P=40).

**Supplemental Figure 3.** Genomic distribution and characteristics of predicted regulatory variants. (A) Genomic distribution of high-impact variants (top 10% by effect score). The stacked bar chart shows the percentage distribution across different genomic features, including promoter regions, UTRs, exons, introns, and intergenic regions. (B) Relationship between variant effect scores and minor allele frequency (MAF). The plot shows decreasing mean MAF with increasing effect scores (binned by log10 scale), suggesting stronger selective constraints on high-impact variants. The dashed line indicates the genome-wide average MAF. (C) Example of a functional variant affecting PR1 gene (AT2G14610) expression. Top: gene structure showing the location of SNP chr2:6321143 (T/A). Bottom: Box plot showing differential gene expression between T (*n*=30) and A (*n*=590) alleles (FDR = 6.92E-4). (D) Differential binding predictions for WRKY transcription factors at the PR1 variant site. Bar plots show binding scores for reference (Ref) and alternate (Alt) alleles across four WRKY family members, demonstrating consistently reduced binding affinity with the alternate allele.

**Supplemental Figure 4.** Relationship between transfer learning improvement and training data size in Arabidopsis and wheat. (A) Correlation analysis between performance improvement and number of binding sites in *Arabidopsis*. The y-axis shows the PRAUC difference between deepTFBS-TL and deepTFBS-ST, while the x-axis shows the log10-transformed number of binding sites. A weak negative correlation (R = −0.13, *P*-value = 0.0024) indicates that transfer learning benefits are slightly more pronounced for TFs with fewer binding sites. (B) Similar correlation analysis for wheat TFs, showing a comparable trend (R = −0.15, *P*-value = 0.1136). The weaker correlation and higher P-value likely reflect the smaller number of TFs available for wheat analysis. In both panels, each point represents one TF, and the blue line indicates the linear regression fit. The negative slopes suggest that transfer learning provides greater benefits when training data is limited, though the relationship is modest.

**Supplemental Figure 5.** Quality assessment of WUS DAP-seq data in *Arabidopsis*. (A) Correlation matrix showing the Pearson correlation coefficients between input control and two biological replicates (Ath_rep1 and Ath_rep2). (B) Venn diagram showing the overlap of peaks identified in the two biological replicates. A substantial number of peaks (6,926) were shared between replicates, demonstrating consistency in peak calling. (C-D) Meta-profiles showing the distribution of peak frequencies relative to transcription start sites (TSS) and transcription termination sites (TTS) for replicate 1 (C) and replicate 2 (D). Gray shading indicates 95% confidence intervals. (E) Genomic distribution of WUS binding sites across different functional regions for both replicates. The stacked bar charts show similar distribution patterns between replicates, with predominant binding in promoter and distal intergenic regions.

**Supplemental Figure 6.** Quality assessment of WUS DAP-seq data in maize. (A) Correlation matrix showing the Pearson correlation coefficients between input control and two biological replicates (Zma_rep1 and Zma_rep2). The high correlation between replicates (r = 0.86) indicates good reproducibility. (B) Venn diagram showing the overlap of peaks identified in the two biological replicates. A substantial number of peaks (4,219) were shared between replicates, demonstrating consistency in peak calling. (C-D) Meta-profiles showing the distribution of peak frequencies relative to transcription start sites (TSS) and transcription termination sites (TTS) for replicate 1 (C) and replicate 2 (D). Gray shading indicates 95% confidence intervals. (E) Genomic distribution of WUS binding sites across different functional regions for both replicates. The stacked bar charts show similar distribution patterns between replicates, with predominant binding in promoter and distal intergenic regions.

**Supplemental Figure S7.** Experimental validation of PWM-predicted WUS binding sites using yeast one-hybrid (Y1H) assays.

**Supplemental Figure S8.** Semantic similarity analysis of GO terms enriched in conserved WUS targets.

**Supplemental Figure S9.** Overview of the deepTFBS web server interface and functionality. (A) The database component allows users to query and retrieve predicted binding sites for 512 Arabidopsis and 110 wheat transcription factors. (B) The predictor interface enables users to input DNA sequences for TFBS prediction. Results are presented through an interactive graphical interface highlighting predicted binding sites.

**Supplemental Table S1**. List of 359 *Arabidopsis* transcription factors used for training deepTFBS-MT model.

**Supplemental Table S2**. Performance comparison of different methods (PWM, DeepSEA, DanQ, and deepTFBS-MT) using AUC and PRAUC metrics.

**Supplemental Table S3**. Performance evaluation of deepTFBS-TL model across different transcription factors in *Arabidopsis*.

**Supplemental Table S4**. Cross-species prediction performance of deepTFBS-TL model for 110 wheat transcription factors.

**Supplemental Table S5**. Randomly selected deepTFBS-predicted and PWM-predicted targets using yeast one-hybrid assays.

**Supplemental Table S6**. Genome-wide WUS binding site predictions and their annotations in *Arabidopsis*.

**Supplemental Table S7**. Genome-wide WUS binding site predictions and their annotations in maize.

**Supplemental Table S8**. Genome-wide WUS binding site predictions and their annotations in wheat.

