## Supplemental Figures for "deepTFBS: Improving within- and cross-species prediction of transcription factor binding using deep multi-task and transfer learning"

**Supplemental Figure 1**

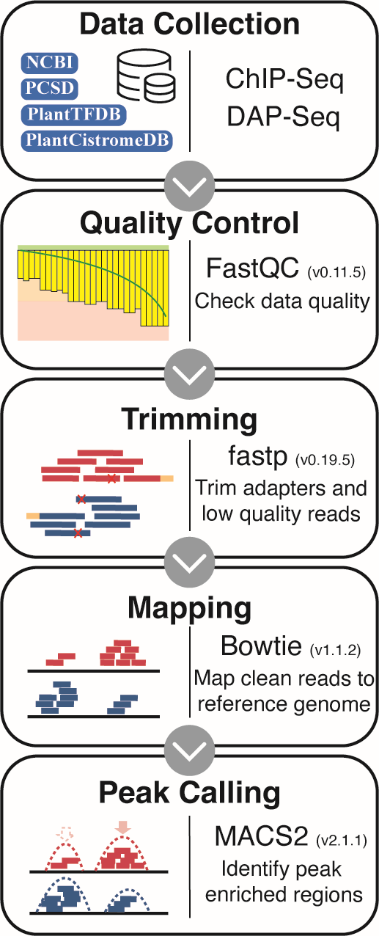

**Supplemental Figure 1.** Standardized computational pipeline for processing DAP-Seq and ChIP-Seq data.

**Supplemental Figure 2**

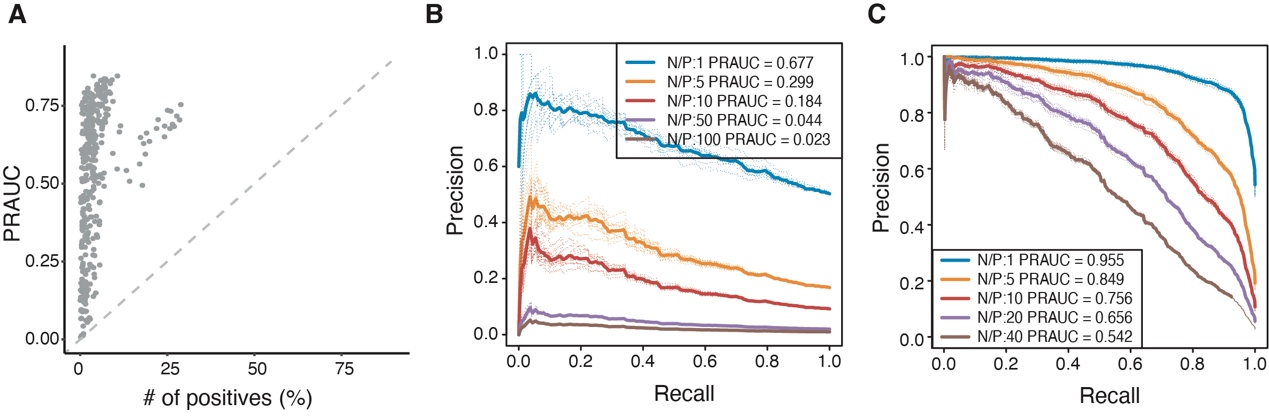

**Supplemental Figure 2. Impact of class imbalance on deepTFBS-MT performance. (**A) Relationship between model performance (PRAUC) and proportion of positive samples in the test dataset for 359 Arabidopsis TFs. Each point represents one TF, showing lower PRAUC values generally correspond to TFs with fewer positive samples. (B) Precision-recall curves for RGA (AT2G01570) at different negative-to-positive (N/P) ratios. The model’s performance decreases significantly as class imbalance increases, with PRAUC dropping from 0.677 (N/P=1) to 0.023 (N/P=100). Dotted lines represent individual runs; solid lines show the average. (C) Precision-recall curves for ATNAC6 (AT5G39610) across varying N/P ratios, demonstrating similar trend of performance decline with increasing class imbalance. PRAUC decreases from 0.955 (N/P=1) to 0.542 (N/P=40).

**Supplemental Figure 3**

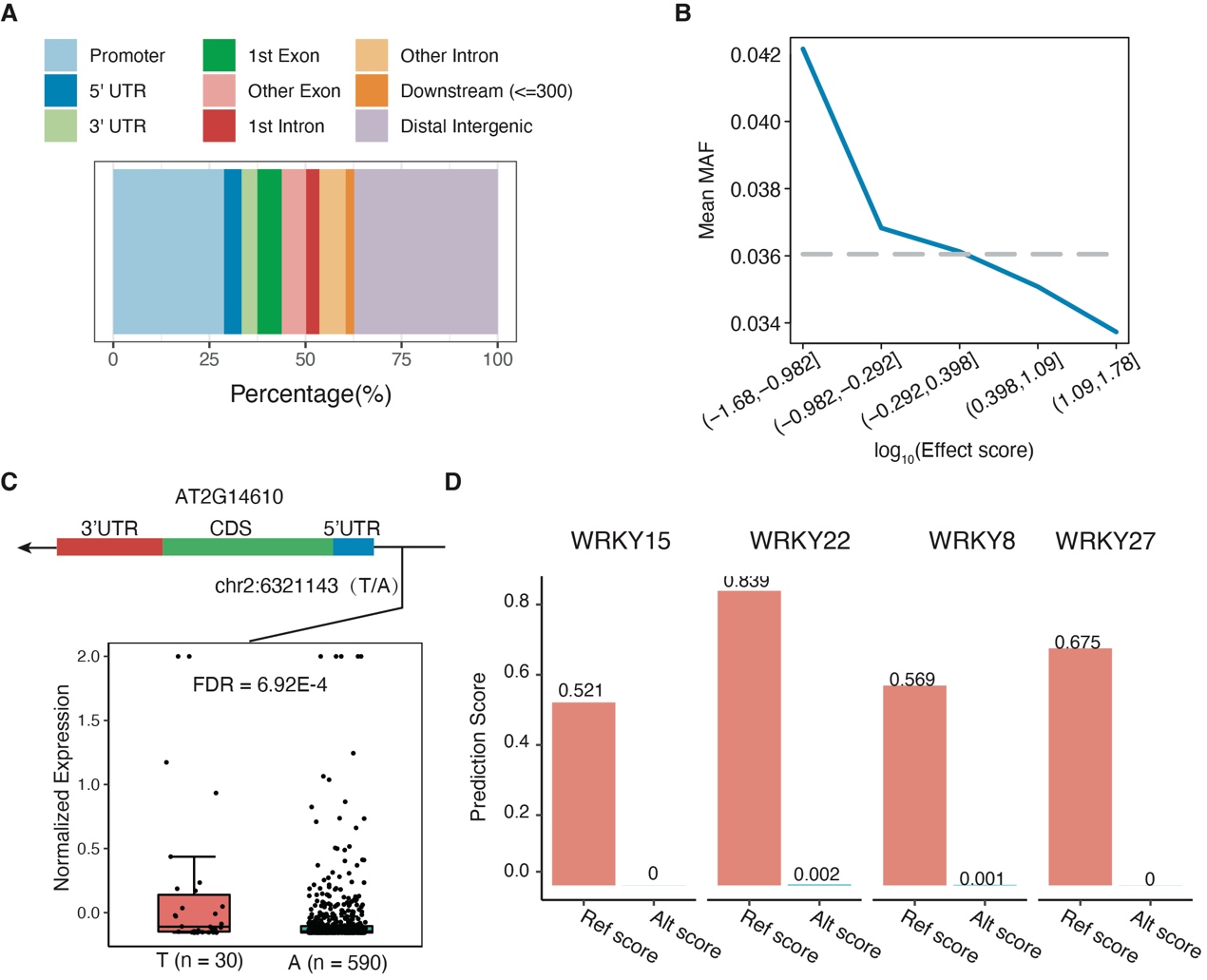

**Supplemental Figure 3. Genomic distribution and characteristics of predicted regulatory variants.** (A) Genomic distribution of high-impact variants (top 10% by effect score). The stacked bar chart shows the percentage distribution across different genomic features, including promoter regions, UTRs, exons, introns, and intergenic regions. (B) Relationship between variant effect scores and minor allele frequency (MAF). The plot shows decreasing mean MAF with increasing effect scores (binned by log10 scale), suggesting stronger selective constraints on high-impact variants. The dashed line indicates the genome-wide average MAF. (C) Example of a functional variant affecting PR1 gene (AT2G14610) expression. Top: gene structure showing the location of SNP chr2:6321143 (T/A). Bottom: Box plot showing differential gene expression between T (n=30) and A (n=590) alleles (FDR = 6.92E-4). (D) Differential binding predictions for WRKY transcription factors at the PR1 variant site. Bar plots show binding scores for reference (Ref) and alternate (Alt) alleles across four WRKY family members, demonstrating consistently reduced binding affinity with the alternate allele.

**Supplemental Figure 4**

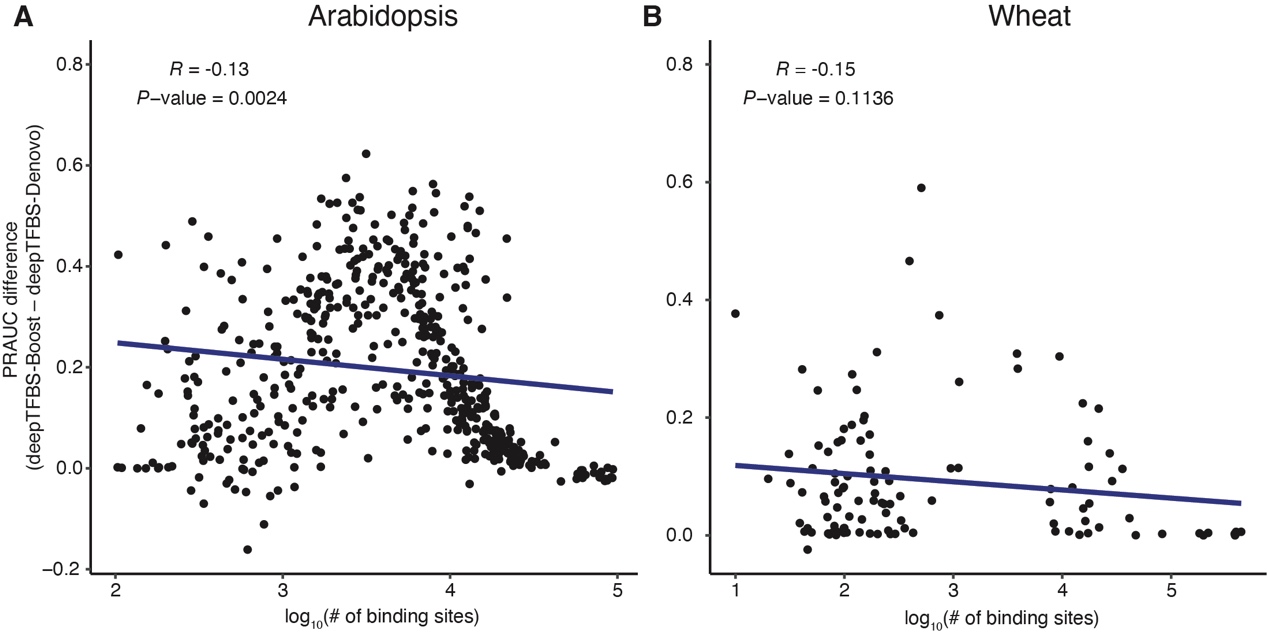

**Supplemental Figure 4. Relationship between transfer learning improvement and training data size in Arabidopsis and wheat.** (A) Correlation analysis between performance improvement and number of binding sites in Arabidopsis. The *y*-axis shows the PRAUC difference between deepTFBS-TL and deepTFBS-ST, while the *x*-axis shows the log10-transformed number of binding sites. A weak negative correlation (R = -0.13, P-value = 0.0024) indicates that transfer learning benefits are slightly more pronounced for TFs with fewer binding sites. (B) Similar correlation analysis for wheat TFs, showing a comparable trend (R = -0.15, *P*-value = 0.1136). The weaker correlation and higher *P*-value likely reflect the smaller number of TFs available for wheat analysis. In both panels, each point represents one TF, and the blue line indicates the linear regression fit. The negative slopes suggest that transfer learning provides greater benefits when training data is limited, though the relationship is modest.

**Supplemental Figure 5**

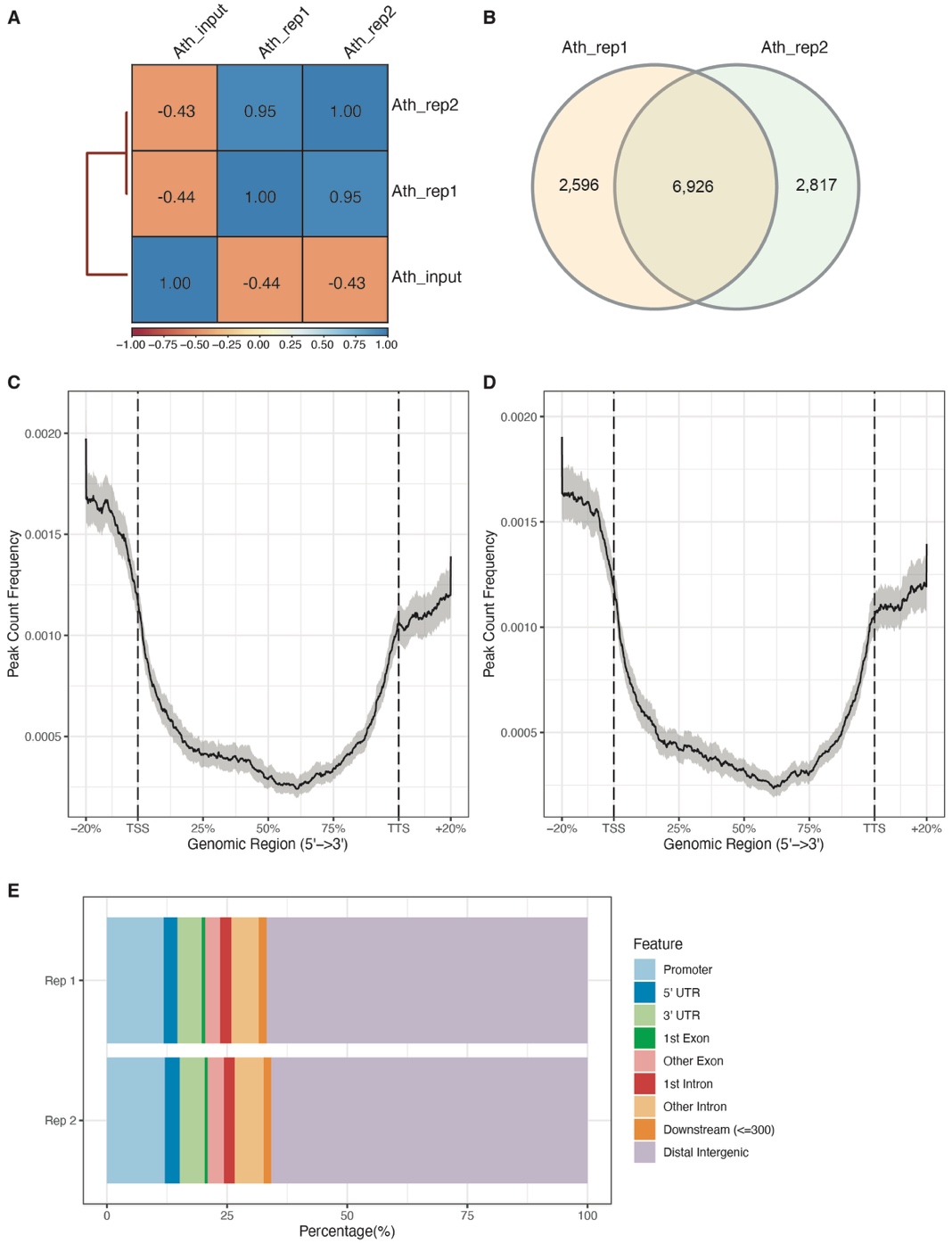

**Supplemental Figure 5. Quality assessment of WUS DAP-seq data in Arabidopsis.** (A) Correlation matrix showing the Pearson correlation coefficients between input control and two biological replicates (Ath_rep1 and Ath_rep2). (B) Venn diagram showing the overlap of peaks identified in the two biological replicates. A substantial number of peaks (6,926) were shared between replicates, demonstrating consistency in peak calling. (C-D) Meta-profiles showing the distribution of peak frequencies relative to transcription start sites (TSS) and transcription termination sites (TTS) for replicate 1 (C) and replicate 2 (D). Gray shading indicates 95% confidence intervals. (E) Genomic distribution of WUS binding sites across different functional regions for both replicates. The stacked bar charts show similar distribution patterns between replicates, with predominant binding in promoter and distal intergenic regions.

**Supplemental Figure 6**

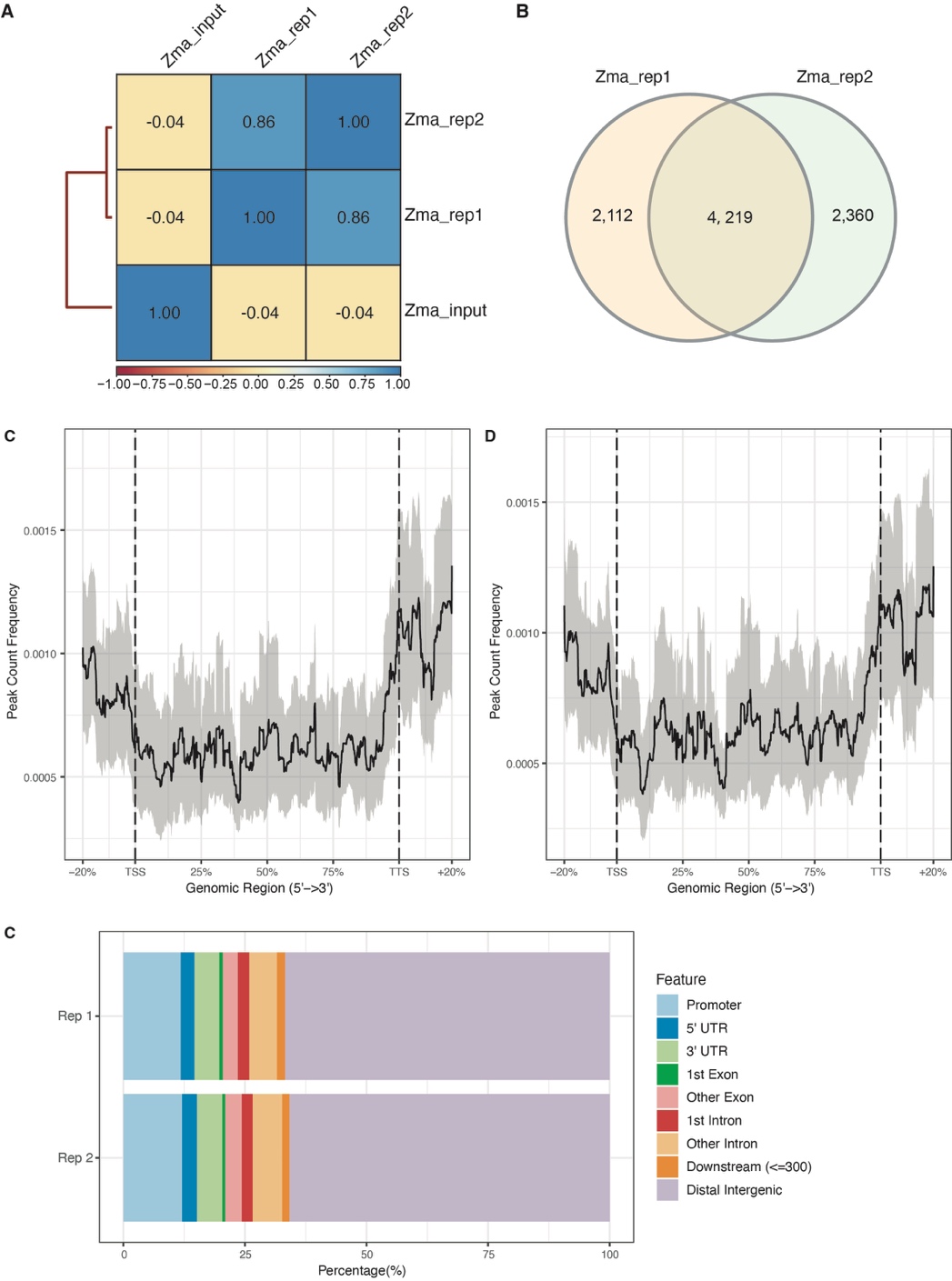

**Supplemental Figure 6. Quality assessment of WUS DAP-seq data in maize.** (A) Correlation matrix showing the Pearson correlation coefficients between input control and two biological replicates (Zma_rep1 and Zma_rep2). The high correlation between replicates (r = 0.86) indicates good reproducibility. (B) Venn diagram showing the overlap of peaks identified in the two biological replicates. A substantial number of peaks (4,219) were shared between replicates, demonstrating consistency in peak calling. (C-D) Meta-profiles showing the distribution of peak frequencies relative to transcription start sites (TSS) and transcription termination sites (TTS) for replicate 1 (C) and replicate 2 (D). Gray shading indicates 95% confidence intervals. (E) Genomic distribution of WUS binding sites across different functional regions for both replicates. The stacked bar charts show similar distribution patterns between replicates, with predominant binding in promoter and distal intergenic regions.

**Supplemental Figure 7**

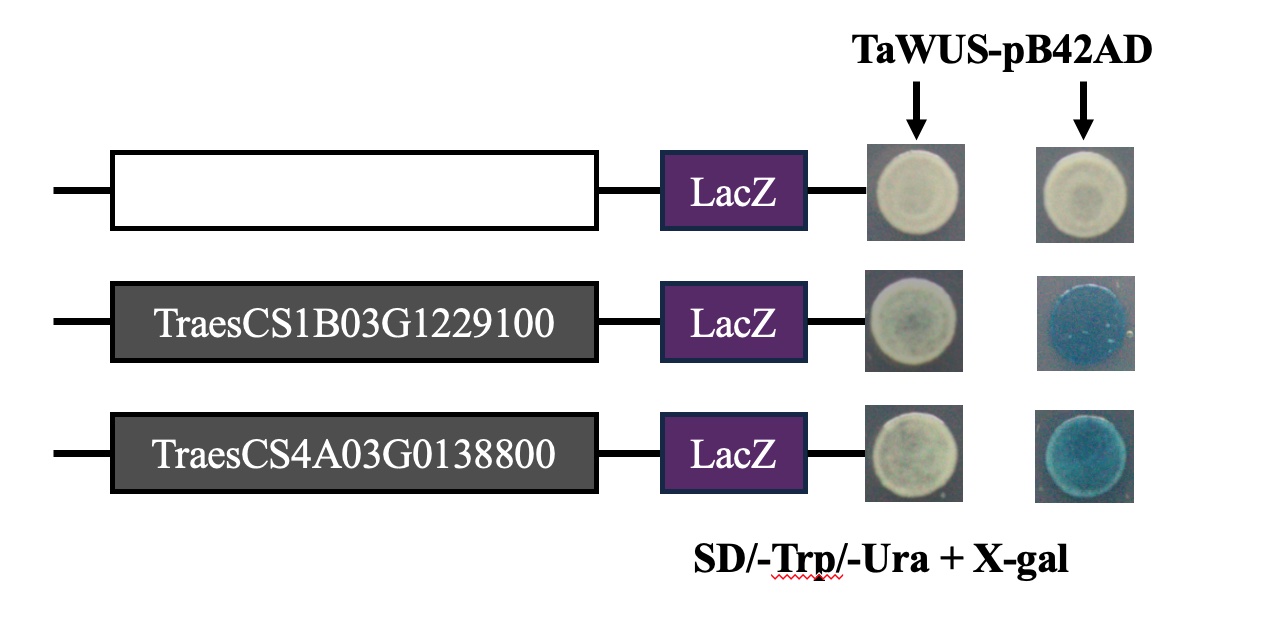

**Supplemental Figure 7. Experimental validation of PWM-predicted WUS binding sites using yeast one-hybrid (Y1H) assays.**

**Supplemental Figure 8**

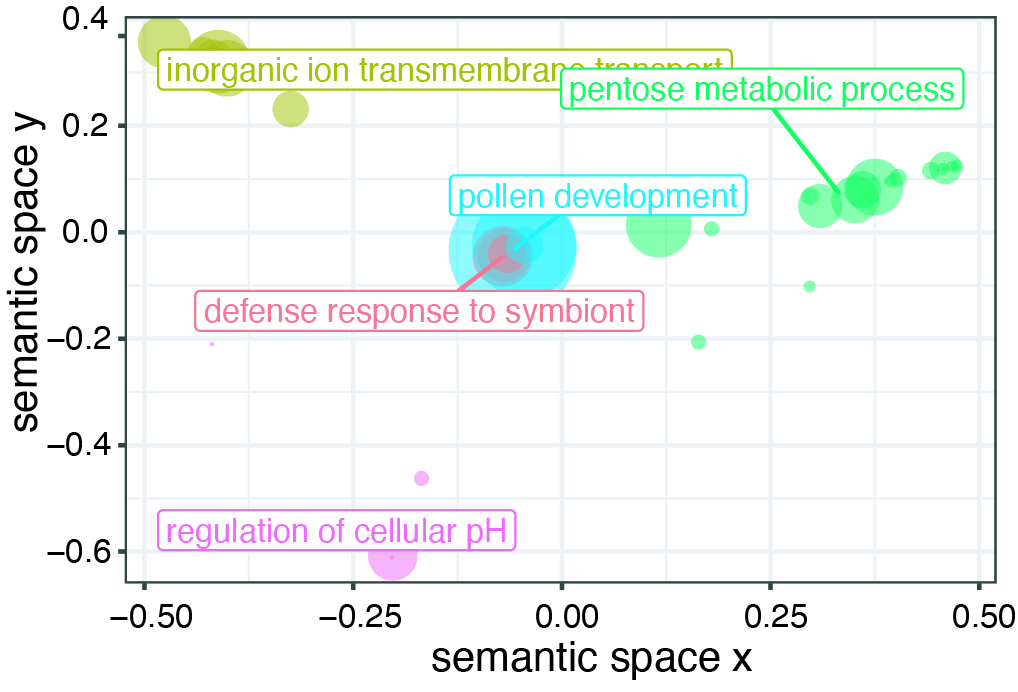

**Supplemental Figure 8. Semantic similarity analysis of GO terms enriched in conserved WUS targets.**

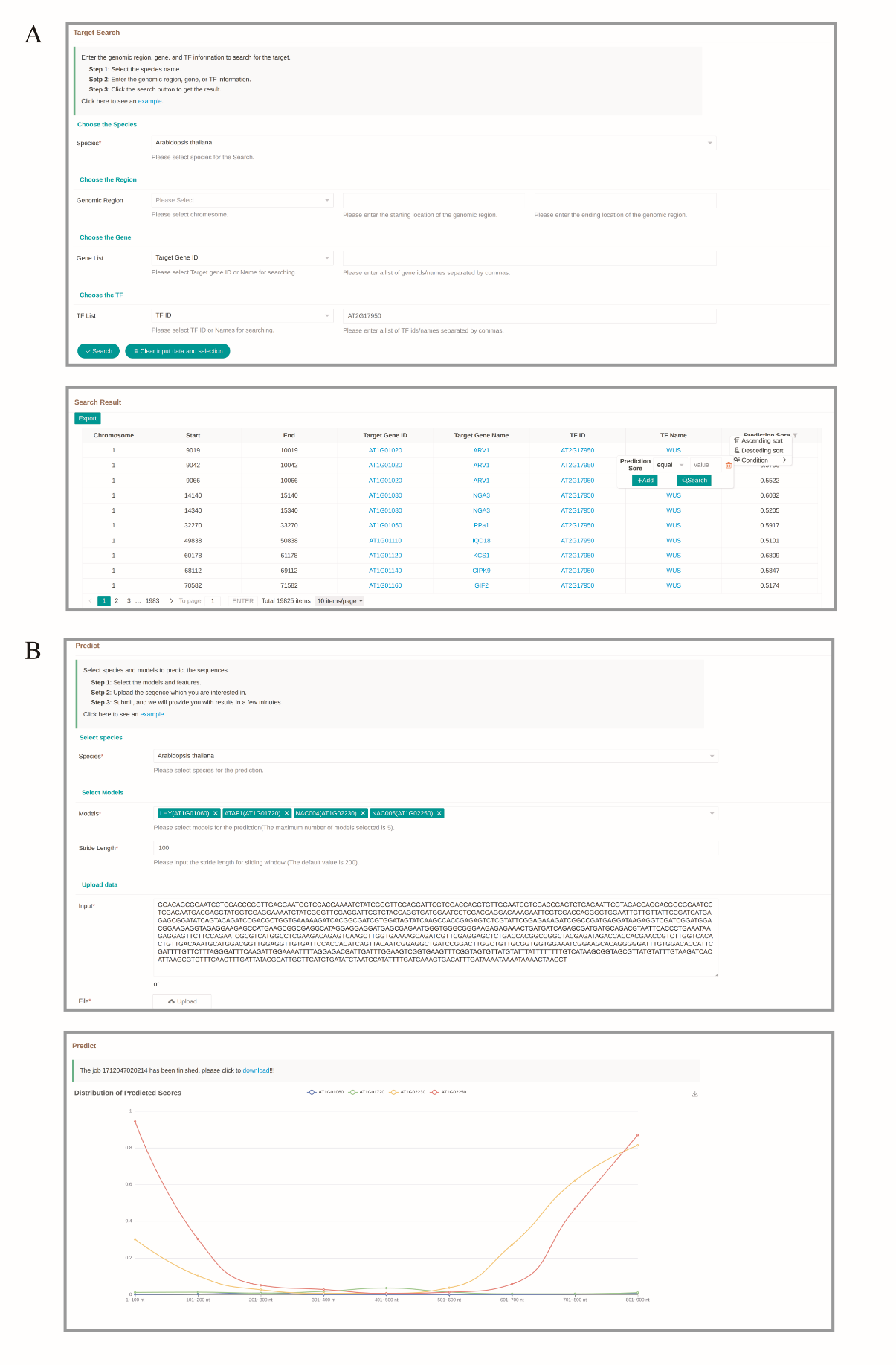

**Supplemental Figure 9. Overview of the deepTFBS web server interface and functionality.** (A) The database component allows users to query and retrieve predicted binding sites for 512 Arabidopsis and 110 wheat transcription factors. (B) The predictor interface enables users to input DNA sequences for TFBS prediction. Results are presented through an interactive graphical interface highlighting predicted binding sites.
